# DeepSCM: an efficient convolutional neural network surrogate model for the screening of therapeutic antibody viscosity

**DOI:** 10.1101/2022.03.12.484110

**Authors:** Pin-Kuang Lai

**Affiliations:** Department of Chemical Engineering and Materials Science, Stevens Institute of Technology, Hoboken, New Jersey 07030

**Author notes:** Corresponding Author: Pin-Kuang Lai.

**Keywords:** deep learning, convolutional neural network, molecular dynamics simulations, spatial charge map, antibody viscosity, developability

## Abstract

Predicting high concentration antibody viscosity is essential for developing subcutaneous administration. Computer simulations provide promising tools to reach this aim. One such model is the spatial charge map (SCM) proposed by Agrawal and coworkers (*mAbs*. **2015**, 8(1):43–48). SCM applies molecular dynamics simulations to calculate a score for the screening of antibody viscosity at high concentrations. However, molecular dynamics simulations are computationally costly and require structural information, a significant application bottleneck. In this work, high throughput computing was performed to calculate the SCM scores for 6596 nonredundant antibody variable regions. A convolutional neural network surrogate model, DeepSCM, requiring only sequence information, was then developed based on this dataset. The linear correlation coefficient of the DeepSCM and SCM scores achieved 0.9 on the test set (N=1320). The DeepSCM model was applied to screen the viscosity of 38 therapeutic antibodies that SCM correctly classified and resulted in only one misclassification. The DeepSCM model will facilitate high concentration antibody viscosity screening. The code and parameters are freely available at https://github.com/Lailabcode/DeepSCM.

## 1. Introduction

Subcutaneous administration of therapeutic antibodies requires low volume and high concentration formulations.^1,2^ At high protein concentrations, some antibodies might exhibit elevated viscosity beyond the syringeability limit.^3^ However, most antibodies have low viscosity at low concentrations.^4^ It is a challenge to identify potential problematic antibodies during discovery. Additionally, there are not enough materials for high concentration measurements in the early-stage development.

Therefore, developing computational tools to assist viscosity screening early is very attractive. There are two types of computational models for predicting antibody viscosity. The first type is based on statistical modeling, and the second is based on physical modeling. Tomer et al. applied regression analysis to develop models to predict concentration-dependent antibody viscosity.^5^ Sharma et al. also proposed a linear model based on three parameters to predict viscosity at 180 mg/ml (pH 5.5 and 200 mM arginine-HCl).^6^ Recently, machine learning has been applied to predict high concentration antibody viscosity.^7–9^ Because of limited experimental data, only conventional machine learning algorithms such as logistic regression, support vector classification, and decision tree classification were applied. The machine learning features relied on domain expertise and published literature. One machine learning model was developed from 27 therapeutic monoclonal antibodies (mAbs) to classify low and high viscosity.^7^ It is a decision tree model with two features, high viscosity index and mAbs net charge. This machine learning model was applied to predict viscosity for 14 immunoglobulins G1 (IgG1) and 14 immunoglobulins G4 S228P (IgG4P) therapeutic mAbs at 150 mg/mL in a subsequent study. The accuracy for IgG1 was 0.86. The accuracy for IgG4P was 0.71. In recent work, this machine learning model was applied to predict antibody viscosity at 150 mg/mL for 20 preclinical/clinical mAbs. The accuracy was 0.55, significantly worse than that of marketed mAbs.

Physical models include molecular dynamics simulations (MD)^10^, coarse-grained (CG) simulations^11–15^, and theoretical models.^16^ One significant advantage of these physical models is that they require little or no training data for prediction. The apparent drawback for MD and CG simulations is the expensive computational time. Spatial charge map (SCM)^10^ was developed, assuming that most antibody regions at formulation conditions carry net positive charges. If there are negative charge patches on the variable fragment (Fv) regions, the molecules tend to self-associate in solution, increasing the solution viscosity. The calculation of the SCM score requires MD simulations. The SCM model was compared with the machine learning model for the 14 IgG1/14 IgG4P commercial mAbs^8^ and 20 preclinical/clinical mAbs^9^ mentioned earlier to evaluate the prediction accuracy. The accuracy for the 14 IgG1 and 14 IgG4 commercial mAbs were 0.93 and 0.79, respectively. The accuracy for the 20 preclinical/clinical mAbs was 0.60. The performance of the SCM model in these two studies was slightly better than that of the machine learning model trained from 27 commercial mAbs. These results indicated that SCM is a reasonable predictor for high concentration viscosity. The SCM score has also been used as a machine learning feature to predict antibody aggregation.^9,17^ The obstacles to applying SCM are the computational cost and difficulties in model construction.

Deep learning is a subset of machine learning. It consists of multi-layer neural networks with many hidden units.^18^ The most common architectures for deep learning are artificial neural networks (ANN), convolutional neural networks (CNN), and recurrent neural networks (RNN). The key difference between deep learning and traditional machine learning is the ability to learn features by itself. Conventional machine learning requires predefined features from human expertise. Therefore, deep learning can learn high-level features from the data and works better with larger datasets. Deep learning has been applied to predict a variety of antibody properties.^19^ For example, DeepH3^20^ and DeepAb^21^ were developed to predict antibody structure. Deep learning was also implemented to predict antibody binders to a target antigen.^22^ Another great application is antibody-specific B-cell epitope prediction by DRREP.^23^ Antibody apparent solubility could also be evaluated by solPredict.^24^ Currently, deep learning has not been applied to predict antibody viscosity due to limited experimental data publicly available. This study aims to apply deep learning to develop a surrogate model for SCM, called DeepSCM. An extensive set of antibody Fv sequences (N=6596) was collected, and their homology models were built to run MD simulations for the SCM scores calculation. The deep learning algorithm used the preprocessed antibody sequences as input and the SCM scores obtained from MD simulations as output for model training. Eventually, an efficient DeepSCM surrogate model was developed based on the CNN architecture. DeepSCM will facilitate antibody developability screening in the early-stage design.

## 2. Materials and Methods

### 2.1 Antibody sequence datasets

Antibody sequences were retrieved from SAbDab^25^, a curated dataset of all antibody structures in the Protein Data Bank and AbYsis,^26^ a web-based database of antibody sequence and structure data. Only those sequences with paired Fv regions were retained. Duplicated antibody sequences were removed.

### 2.2 Preprocessing of antibody sequences

Antibody sequences have variable lengths; however, the input sequence length for the deep learning algorithms should be the same. The same input size was achieved by annotating the antibody sequences in the IMGT numbering scheme using ANARCI.^27^ The heavy chain variable region was from H1 to H128, and the light chain variable region was from L1 to L127. Gaps in the antibody sequences were filled with dashes. Insertion was numbered by appending a capital letter to the corresponding position number.

A few criteria were enforced to select antibody sequences. First, antibody sequences having insertion in the variable regions were removed from the dataset. One exception is the CDRH3 region, which has the highest sequence diversity among the variable regions. The maximum length of the CDRH3 region (H105-H117) allowed in this work was 30. The additional positions are 111A, 111B, 111C, 111D, 111E, 111F, 111G, 111H, 112I, 112H, 112G, 112F, 112E, 112D, 112C, 112B, 112A. The length of the CDRL1 (L27-L38) and the CDRH1 (H27-H38) regions was 12. The length of the CDRL2 (L56-L65) and the CDRH2 (H56-H65) regions was 10. The length of the CDRL3 (L105-L117) regions was 13. Overall, the lengths of the heavy chain variable regions and the light chain variable regions (including gaps) were 145 and 127, respectively. Second, the heavy chain and light chain variable regions should only have two cysteine residues, respectively, on positions 23 and 104. Antibody sequences that did not meet the criteria were removed from the dataset. Third, antibody sequences that failed to generate homology models on the Fv regions were removed from the dataset. These steps resulted in 6596 nonredundant antibody Fv sequences.

### 2.3 Computational Modeling of mAbs

The homology models of the variable regions were generated by ABodyBuilder.^28^ MODELLER was used to run *ab initio* modeling in case CDR templates were not found.^29^ IMGT numbering was used to annotate the final models.

### 2.4 Molecular Dynamics Simulations

Molecular dynamics simulations were performed using all-atom antibody Fv structures with explicit solvent using the TIP3P water model.^30^ Simulation boxes were set up using VMD to place a single antibody Fv structure in a water box extending 12 Å beyond the protein surface.^31^ The salt concentration was 15 mM NaCl. Counterions were added to neutralize the system charge. Simulations were performed at 300 K and 1 atm in the NPT ensemble, using the NAMD software package and the CHARMM36m force field.^32–34^ The histidine residues were protonated. Electrostatic interactions were treated with the Particle Mesh Ewald (PME) method, and van der Waals interactions were calculated using a switching distance of 10 Å and a cutoff of 12 Å.^35^ After energy minimization, the system was gradually heated up from 100 K to 300 K at an interval of 5 K over 200 ps. The heavy atoms were constrained during the heating process. The scaling factor for harmonic constraint energy function was 2.5 kcal/Å^2^. After heating, the constraints were gradually turned off by changing the scaling factor from 2.0, 1.5, 1.0, 0.5 kcal/ Å^2^ over 80 ps. The integration time step was set to 1 fs during heating and relaxation steps. In the end, the production run was 1 ns, and the integration time step was set to 2 fs by applying rigid bond constraints to hydrogen-containing bonds.

### 2.5 Calculation of Spatial Charge Map Scores

The spatial charge map (SCM) is a score to rank antibodies that exhibit high viscosity in a condensed protein solution. The calculation of SCM scores follows previous work.^10^ Briefly, the atomic SCM value has the following form.

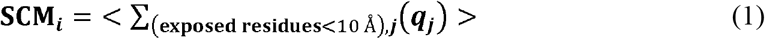

where <> indicates ensemble average from MD simulations. The atomic SCM value is the summation of all the partial charges (*q*_*j*_) on the surrounding atom *j*, which belongs to exposed residues whose side-chain atoms are within 10 Å of atom *i*. The exposed residues are defined as the solvent-accessible surface area of the side-chain atoms ≥10 Å^2^. Partial charges were taken from the forcefield for MD simulations. The SCM score on the Fv region is expressed as

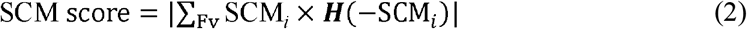

where *H* is the Heaviside function, and |. | is the absolute value function.

### 2.6 Machine and deep learning model construction

Machine learning models were built in Python 3.7.12. The train_test_split function was implemented using Scikit-learn v1.0.2.^36^ CNNs were built using the Keras v2.7.0 Sequential model^37^ as a wrapper for TensorFlow v2.7.0.^38^ The CNN architecture and hyperparameters were optimized by performing different combinations of CNN layers (1, 2, 3), numbers of filters (32, 64, 128, 256) and kernel sizes (3, 5, 7). The MAE values of the best validation model were used to evaluate the model performance.

### 2.7 Machine and deep learning model training and testing

The dataset used for regression was split into a training set (60%), a validation set (20%), and a test set (20%). For the CNN model, the activation function was Relu. Figure 3 (A) illustrates the best architecture and hyperparameters. During model training, the number of epochs was 50, the batch size was 64, the optimizer was Adam^39^, and the loss function was MAE. The best models were recorded by ModelCheckpoint from keras.callbacks. The CNN architecture and weights were saved to JSON and HDF5 (H5) formats, respectively.

### 2.8 Sequenced-based viscosity prediction models

The Sharma method is based on the Fv charge, the product of VH and VL charges, and the hydrophobicity index at pH 6.0 to predict viscosity at 180 mg/mL using only VH and VL sequences.^6^ The TAP: Therapeutic Antibody Profiler web server was utilized to predict the number of developability flags.^40^

## 3. Results

### 3.1 Comparison of the viscosity prediction models for preclinical to marketed mAbs

Before developing a surrogate model for the structure-based SCM model, it is imperative to compare the accuracy of the SCM model with other sequence-based viscosity prediction models. Table 1 compares the SCM model, the Sharma method^6^, and the therapeutic antibody profiler (TAP).^40^ The Sharma model predicts viscosity at 180 mg/mL, is based on the Fv charge, the product of heavy chain variable region (VH) and light chain variable region (VL) charges, and the hydrophobicity index at a given pH. In table 1, the predicted viscosity was linearly interpolated to 150 mg/mL. TAP is a general antibody developability predictor based on five metrics. The 61 mAbs viscosity data at 150 mg/mL were from our previous works.^7–9^ The first 20 data were preclinical and clinical stage mAbs, and the remaining 41 were commercial mAbs. The solution was at pH 5.5 to pH 6.0 in a 10-20 histidine-HCl buffer. There were 14 high viscosity mAbs (>30 cP). For prediction models, the cutoff value for the SCM score is 1000, and the TAP criterion is with at least one flag. The prediction accuracy for the SCM model, Sharma method, and TAP were 0.70, 0.61, and 0.64, respectively. For preclinical and clinical mAbs only, the accuracy for the three models were 0.70, 0.55, and 0.60, respectively. The SCM model performed better than the other two sequence-based models, and the prediction results were consistent for the preclinical/clinical and the commercial mAbs. It is noted that the Sharma model was fitted with the viscosity data in a buffered solution at pH 5.5 and 200 mM arginine-HCl. The difference between the solution conditions may affect the performance of the model. Additionally, the TAP flag may indicate other developability issues other than high viscosity. Overall, the SCM model, although considering only negative surface charges and having room for improvement, is currently a good viscosity prediction model. The implementation bottleneck is the computationally expensive MD simulation. Therefore, developing a machine learning surrogate model for the SCM calculation is preferred.

**Table 1.** Summary of the experimental viscosity at 150 mg/mL and viscosity prediction for 61 preclinical to commercial mAbs. The SCM score is from the MD simulation, and the Sharma model is a viscosity regression model. TAP is the therapeutic antibody profiler. The red labels indicate predicted or experimental high viscosity. The cutoff value for high viscosity is 30 cP for the experiment and the Sharma model. The cutoff value for the SCM score is 1000. The condition for TAP is with at least one flag. Their names in the source data are also listed.

| Source | Experiment | Prediction |  |  | Accuracy |  |  |
| --- | --- | --- | --- | --- | --- | --- | --- |
|  | Viscosity<br>(cP)<br>at 150<br>mg/mL | SCM<br>score | Sharma<br>model<br>(cP) | TAP<br>flags | SCM<br>score | Sharma<br>model<br>(cP) | TAP<br>flags |
| mAb1<sup>9</sup> | 16.6 | 673.3 | 17.3 | 1 | 1 | 1 | 0 |
| mAb2<sup>9</sup> | 6.5 | 1173.9 | 31.7 | 0 | 0 | 0 | 1 |
| mAb3<sup>9</sup> | 7.3 | 905.2 | 16.4 | 1 | 1 | 1 | 0 |
| mAb4<sup>9</sup> | 9.7 | 759.3 | 10.9 | 0 | 1 | 1 | 1 |
| mAb5<sup>9</sup> | 7.0 | 831.6 | 14.3 | 1 | 1 | 1 | 0 |
| mAb6<sup>9</sup> | 10.4 | 609.0 | 13.7 | 0 | 1 | 1 | 1 |
| mAb7<sup>9</sup> | 23.3 | 1025.6 | 42.0 | 0 | 0 | 0 | 1 |
| mAb8<sup>9</sup> | 16.1 | 2273.6 | 40.5 | 1 | 0 | 0 | 0 |
| mAb9<sup>9</sup> | 6.2 | 708.6 | 70.8 | 0 | 1 | 0 | 1 |
| mAb10<sup>9</sup> | 227.5 | 798.6 | 1.7 | 1 | 0 | 0 | 1 |
| mAb11<sup>9</sup> | 26.0 | 969.7 | 23.4 | 2 | 1 | 1 | 0 |
| mAb12<sup>9</sup> | 108.3 | 923.9 | 22.8 | 1 | 0 | 0 | 1 |
| mAb13<sup>9</sup> | 93.0 | 1118.4 | 35.9 | 1 | 1 | 1 | 1 |
| mAb14<sup>9</sup> | 102.5 | 1163.5 | 36.1 | 2 | 1 | 1 | 1 |
| mAb15<sup>9</sup> | 21.3 | 959.4 | 44.9 | 2 | 1 | 0 | 0 |
| mAb16<sup>9</sup> | 115.6 | 1256.5 | 5.6 | 2 | 1 | 0 | 1 |
| mAb17<sup>9</sup> | 13.1 | 820.3 | 13.0 | 1 | 1 | 1 | 0 |
| mAb18<sup>9</sup> | 13.6 | 978.5 | 45.5 | 0 | 1 | 0 | 1 |
| mAb19<sup>9</sup> | 7.8 | 1644.6 | 11.7 | 1 | 0 | 1 | 0 |
| mAb20<sup>9</sup> | 48.9 | 1156.0 | 48.1 | 1 | 1 | 1 | 1 |
| mAb1<sup>7</sup> | 14.4 | 1401.8 | 32.9 | 1 | 0 | 0 | 0 |
| mAb2<sup>7</sup> | 20.9 | 763.4 | 36.9 | 0 | 1 | 0 | 1 |
| mAb3<sup>7</sup> | 14.9 | 989.4 | 37.4 | 0 | 1 | 0 | 1 |
| mAb4<sup>7</sup> | 93.4 | 1234.7 | 18.3 | 0 | 1 | 0 | 0 |
| mAb5<sup>7</sup> | 8.6 | 1152.4 | 41.1 | 0 | 0 | 0 | 1 |
| mAb6<sup>7</sup> | 9.0 | 651.0 | 21.8 | 0 | 1 | 1 | 1 |
| mAb7<sup>7</sup> | 29.0 | 1275.6 | 58.7 | 0 | 0 | 0 | 1 |
| mAb8<sup>7</sup> | 12.9 | 1132.5 | 43.6 | 0 | 0 | 0 | 1 |
| mAb9<sup>7</sup> | 52.3 | 950.6 | 22.7 | 0 | 0 | 0 | 0 |
| mAb10<sup>7</sup> | 10.2 | 763.8 | 6.2 | 0 | 1 | 1 | 1 |
| mAb11<sup>7</sup> | 100.3 | 947.4 | 26.2 | 0 | 0 | 0 | 0 |
| mAb12<sup>7</sup> | 7.5 | 814.2 | 13.7 | 0 | 1 | 1 | 1 |
| mAb13<sup>7</sup> | 12.5 | 995.7 | 7.7 | 1 | 1 | 1 | 0 |
| mAb14<sup>7</sup> | 23.4 | 793.1 | 29.9 | 0 | 1 | 1 | 1 |
| mAb15<sup>7</sup> | 12.9 | 669.3 | 10.8 | 0 | 1 | 1 | 1 |
| mAb16<sup>7</sup> | 10.0 | 1193.9 | 15.8 | 0 | 0 | 1 | 1 |
| mAb17<sup>7</sup> | 210.9 | 1205.4 | 30.8 | 3 | 1 | 1 | 1 |
| mAb18<sup>7</sup> | 7.1 | 706.5 | 14.8 | 1 | 1 | 1 | 0 |
| mAb19<sup>7</sup> | 21.2 | 1065.1 | 46.2 | 1 | 0 | 0 | 0 |
| mAb20<sup>7</sup> | 24.4 | 834.2 | 13.1 | 0 | 1 | 1 | 1 |
| mAb21<sup>7</sup> | 8.6 | 929.5 | 29.1 | 0 | 1 | 1 | 1 |
| mAb22<sup>7</sup> | 9.0 | 1081.1 | 21.1 | 1 | 0 | 1 | 0 |
| mAb23<sup>7</sup> | 22.9 | 432.0 | 5.9 | 0 | 1 | 1 | 1 |
| mAb24<sup>7</sup> | 90.5 | 1081.7 | 50.4 | 0 | 1 | 1 | 0 |
| mAb25<sup>7</sup> | 7.4 | 617.1 | 2.0 | 0 | 1 | 1 | 1 |
| mAb26<sup>7</sup> | 10.3 | 612.8 | 12.9 | 0 | 1 | 1 | 1 |
| mAb27<sup>7</sup> | 103.8 | 942.7 | 28.9 | 0 | 0 | 0 | 0 |
| Adalimumab IgG1<sup>8</sup> | 12.8 | 1401.8 | 33.1 | 1 | 0 | 0 | 0 |
| Atezolizumab IgG1<sup>8</sup> | 22.3 | 673.7 | 35.3 | 0 | 1 | 0 | 1 |
| Basiliximab IgG1<sup>8</sup> | 8.6 | 561.4 | 3.0 | 1 | 1 | 1 | 0 |
| Bevacizumab IgG1<sup>8</sup> | 6.7 | 989.4 | 37.4 | 0 | 1 | 0 | 1 |
| Cetuximab IgG1<sup>8</sup> | 55.7 | 1234.7 | 18.3 | 0 | 1 | 0 | 0 |
| Ganitumab IgG1<sup>8</sup> | 10.9 | 711.9 | 12.0 | 0 | 1 | 1 | 1 |
| Golimumab IgG1<sup>8</sup> | 7.6 | 763.8 | 6.2 | 0 | 1 | 1 | 1 |
| Ipilimumab IgG1<sup>8</sup> | 18.0 | 773.5 | 6.9 | 0 | 1 | 1 | 1 |
| Natalizumab IgG1<sup>8</sup> | 11.3 | 1067.8 | 10.7 | 0 | 0 | 1 | 1 |
| Omalizumab IgG1<sup>8</sup> | 60.0 | 1205.4 | 30.8 | 3 | 1 | 1 | 1 |
| TGN1412 IgG1<sup>8</sup> | 6.1 | 684.8 | 9.8 | 1 | 1 | 1 | 0 |
| Trastuzumab IgG1<sup>8</sup> | 9.3 | 612.8 | 12.9 | 0 | 1 | 1 | 1 |
| Tremelimumab IgG1<sup>8</sup> | 14.2 | 644.3 | 6.6 | 0 | 1 | 1 | 1 |
| Vesencumab IgG1<sup>8</sup> | 12.0 | 704.2 | 21.1 | 0 | 1 | 1 | 1 |
|  |  |  |  | Accuracy | 0.70 | 0.61 | 0.64 |

### 3.2 Antibody sequence dataset and statistical analysis

The antibody variable region sequences were retrieved from SAbDab and AbYsis databases. After removing redundant sequences and filtering out sequences based on some criteria such as complementarity determining region (CDR) length, the number of cysteine residues, and insertion (detailed in the Materials and Methods section), there were 6596 antibody Fv sequences in the dataset for this study. The Fv sequences covered therapeutic antibodies and nontherapeutic antibodies to increase sequence diversity.

Figure 1 shows the length distribution of different antibody regions in the dataset. The VH length was approximately normal distributed, centered at 119. The number of antibodies having VL length from 106 to 112 was on average 500, except a high peak at 107. The first complementarity determining region of the heavy chain (CDRH1) length had the highest peak at 8. The first complementarity determining region of the light chain (CDRL1) length had the highest peak at 6, and the number of antibodies having a length of 5 and from 7 to 12 was on average 500. For the second complementarity determining region of the heavy chain (CDRH2) length, the highest peak was at 8, and the second-highest peak was at 7. The second complementarity determining region of the light chain (CDRL2) length had the highest peak at 3. For the third complementarity determining region of the light chain (CDRL3) length, the highest peak was at 9. The third complementarity determining region of the heavy chain (CDRH3) length had a wide distribution centered at 12, and there was a long tail extending to 30.

**Figure 1.**
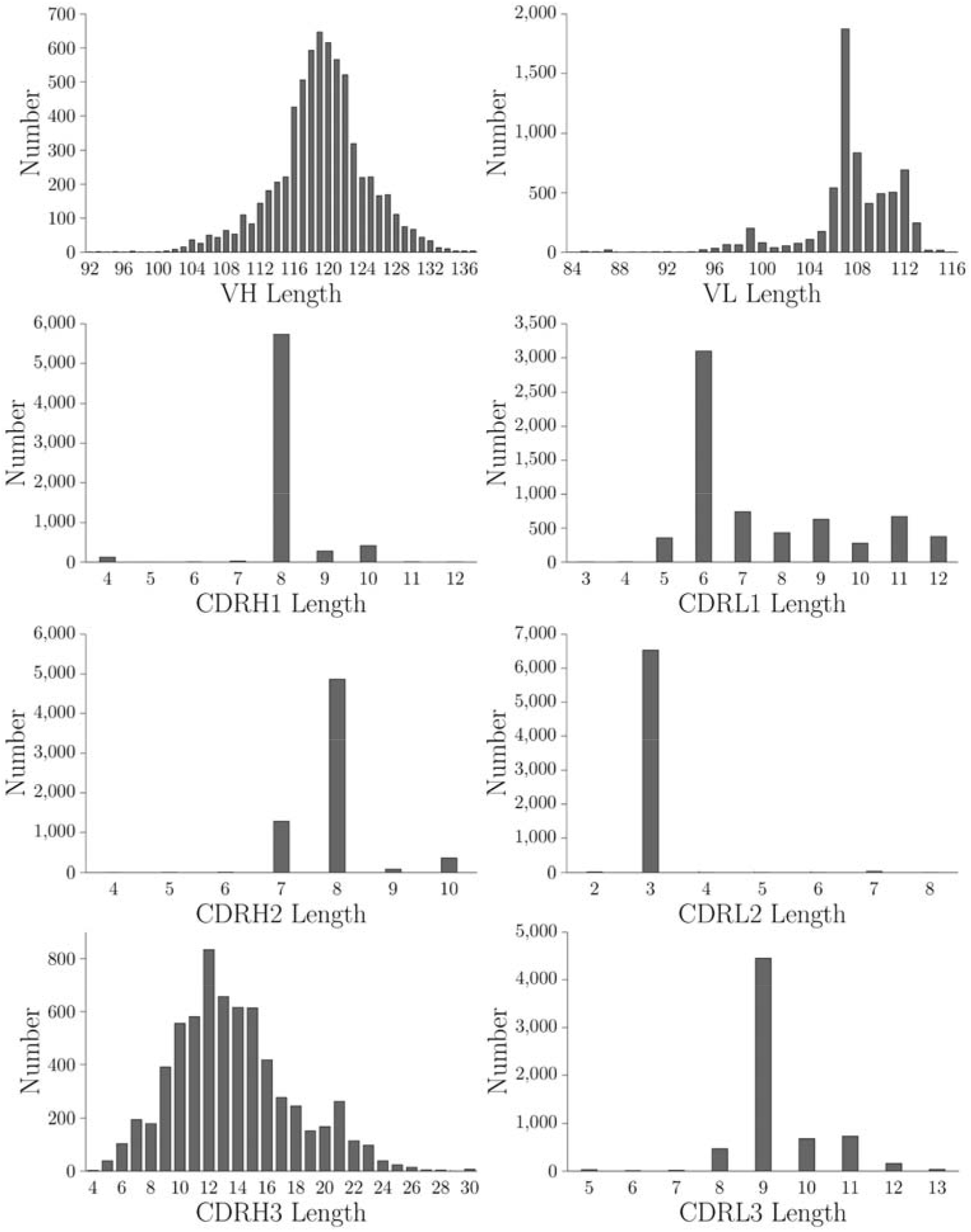
Distribution of VH, VL, CDRH1, CDRH2, CDRH3, CDRL1, CDRL2, and CDRL3 lengths of the 6596 Fv sequences in this study. The CDR regions are based on the IMGT definition.

For the 6596 antibody Fv sequences, there were 21750310 pairwise alignments. Ninety-three percent of pairs have sequence similarities less than 5%. Five percent of pairs have sequence similarity between 5% to 10%, indicating that the antibody dataset has a high sequence diversity. Among the 6596 antibody sequences, 2212 came from alpacas, 1740 came from humans, 2017 came from mice, 59 came from pigs, 18 came from rabbits, and 550 came from rhesus.

### 3.3 MD simulations and SCM calculation of the antibody in the dataset

The homology models of the 6596 antibody variable regions were constructed to perform MD simulations. The SCM scores were calculated by the ensemble averages over 1000 ps. Figure 2 (A) reports the box-and-whisker plot of the SCM score. The SCM scores ranged from 255.2 to 2273.6. The first quartile was 675.8, and the third quartile was 1034.4. The medium was 833.5. There were 187 outliers above the upper Whisker, 1572.4.

**Figure 2.**
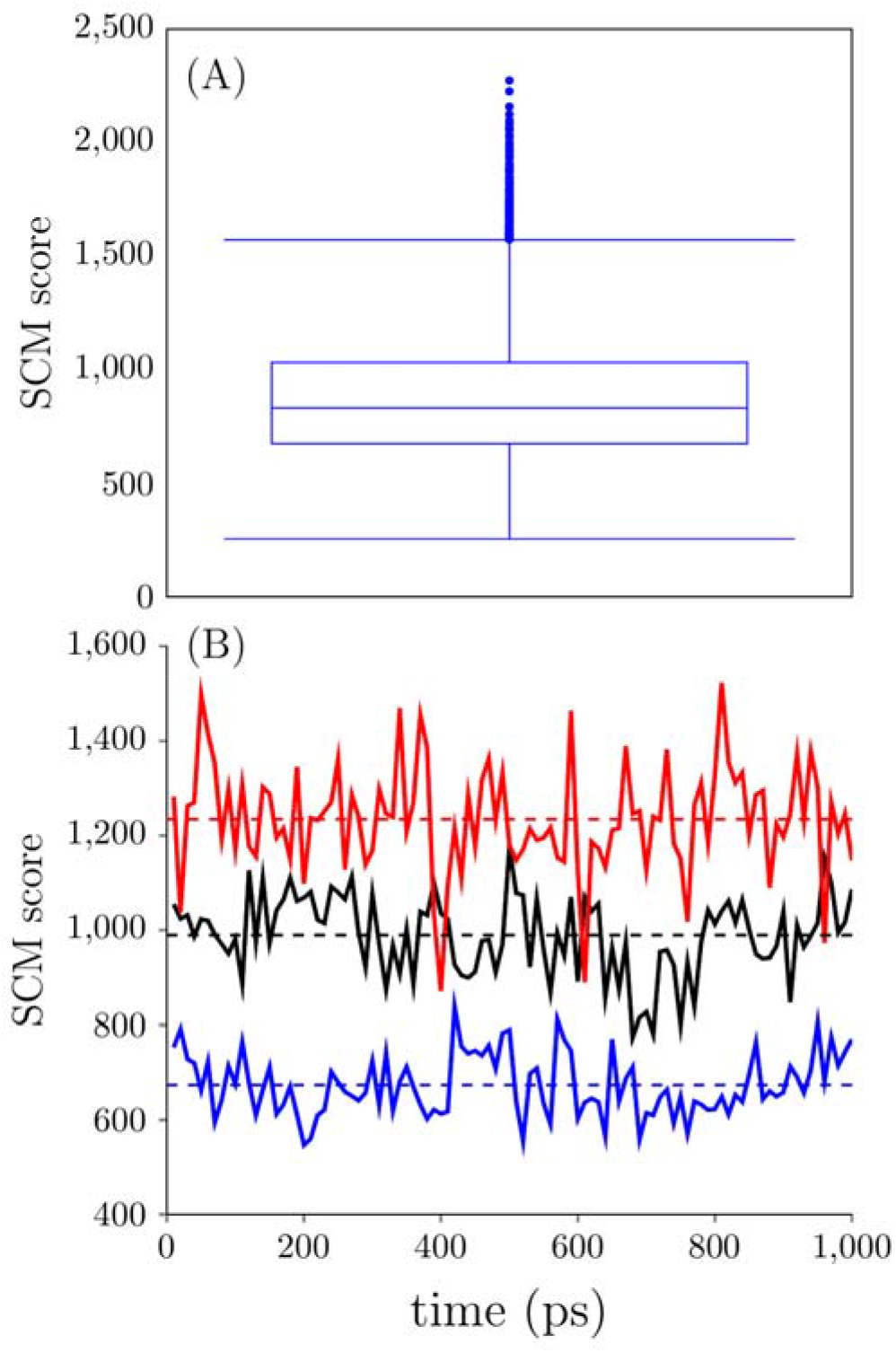
(A) Box-and-Whisker plot of the SCM score (N=6596). (B) Time trajectory for the SCM score of three antibody Fv structures. Their means and standard deviations are 673.7 ± 64.2, 989.3 ± 82.8, and 1234.7 ± 113.7, respectively.

Figure 2 (B) illustrates the time trajectory for three antibodies from low, medium, to high SCM scores. The SCM scores fluctuated around the mean, and the mean converged over the 1000 ps period. A longer simulation time is usually needed for a full-length antibody with a flexible region. However, for a single variable region obtained from homology modeling, the structure is relatively stable, requiring only a short time to equilibrate the system and get converged SCM scores. Therefore, MD simulations using a single variable region are more suitable for high throughput computing for a large antibody dataset.

### 3.4 Antibody sequence preprocessing

Antibody sequences have variable lengths; however, the CNN models require the input to have a fixed size. Therefore, the heavy and light chain variable regions were annotated based on the IMGT numbering scheme. This annotation ensured that the conserved amino acid sequences were aligned. The maximum length of CDRH1 and CDRL1 were both 12, and the total length of CDRH2 and CDRL2 were both 10. The entire length of CDRL3 was 13. For the CDRH3 region, the maximum length in the model was chosen to be 30. All the gaps were padded with dashes. After the preprocessing step, the heavy and light chain variable regions had fixed lengths of 145 and 127, respectively.

### 3.5 CNN model training for the DeepSCM model

CNN models have been shown to perform better than other deep learning models such as ANN and RNN for predicting antibody binders^22^; therefore, the CNN model was chosen for model development in this study. The ratio for training/validation/test split was 60:20:20. The architecture and parameters were optimized by hyperparameter tuning, as shown in Table 2, and Figure 3 (A) illustrates the best CNN architecture and parameters. In the hyperparameter tuning, only the number of 1D CNN layers (1,2,3), number of filters (32,64,128,256), and kernel size (3,5,7) were varied to evaluate the model performance of the best validation model using mean absolute error (MAE).

**Figure 3.**
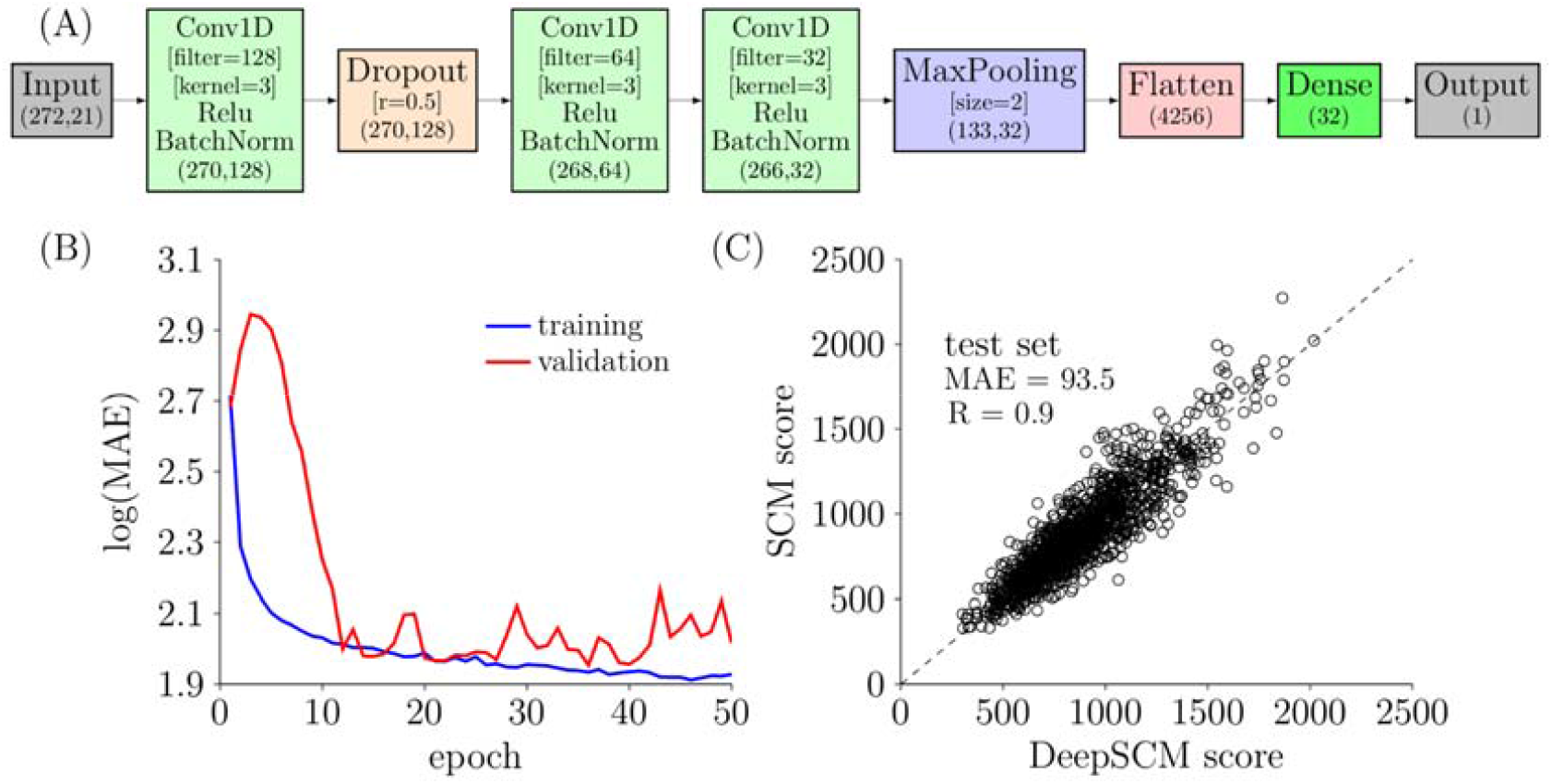
(A) Architecture of the best CNN model. The essential parameters of each layer are shown in the square brackets. The shape of each layer is shown in the round brackets. (B) Loss curves of training and validation loss over epochs. The loss is calculated by mean absolute error (MAE). (C) Scatter plot of the SCM score and the DeepSCM score for the test set. The dashed line is the identity line.

**Table 2.** The MAE value of the best validation model using different numbers of CNN layers and hyperparameters. The optimal architecture and parameters are shown in bold.

| Conv1D_1 |  | Conv1D_2 |  | Conv1D_3 |  | MAE |
| --- | --- | --- | --- | --- | --- | --- |
| filter | kernel | filter | kernel | filter | kernel |  |
| 32 | 3 | - | - | - | - | 214 |
| 32 | 5 | - | - | - | - | 236 |
| 32 | 7 | - | - | - | - | 189 |
| 64 | 3 | - | - | - | - | 307 |
| 64 | 5 | - | - | - | - | 299 |
| 64 | 7 | - | - | - | - | 280 |
| 128 | 3 | - | - | - | - | 332 |
| 128 | 5 | - | - | - | - | 361 |
| 128 | 7 | - | - | - | - | 321 |
| 256 | 3 | - | - | - | - | 402 |
| 256 | 5 | - | - | - | - | 402 |
| 256 | 7 | - | - | - | - | 395 |
| 64 | 3 | 32 | 3 | - | - | 141 |
| 64 | 3 | 32 | 5 | - | - | 159 |
| 64 | 5 | 32 | 3 | - | - | 187 |
| 64 | 5 | 32 | 5 | - | - | 174 |
| 64 | 5 | 32 | 7 | - | - | 142 |
| 64 | 7 | 32 | 5 | - | - | 122 |
| 64 | 7 | 32 | 7 | - | - | 130 |
| 128 | 3 | 64 | 3 | - | - | 146 |
| 128 | 3 | 64 | 5 | - | - | 127 |
| 128 | 5 | 64 | 3 | - | - | 161 |
| 128 | 5 | 64 | 5 | - | - | 143 |
| 128 | 5 | 64 | 7 | - | - | 132 |
| 128 | 7 | 64 | 5 | - | - | 140 |
| 128 | 7 | 64 | 7 | - | - | 121 |
| 256 | 3 | 128 | 3 | - | - | 127 |
| 256 | 3 | 128 | 5 | - | - | 101 |
| 256 | 5 | 128 | 3 | - | - | 153 |
| 256 | 5 | 128 | 5 | - | - | 117 |
| 256 | 5 | 128 | 7 | - | - | 115 |
| 256 | 7 | 128 | 5 | - | - | 123 |
| 256 | 7 | 128 | 7 | - | - | 98 |
| 256 | 3 | 128 | 3 | 64 | 3 | 92 |
| 256 | 3 | 128 | 5 | 64 | 7 | 94 |
| 256 | 5 | 128 | 5 | 64 | 5 | 90 |
| 256 | 7 | 128 | 5 | 64 | 3 | 91 |
| 256 | 7 | 128 | 7 | 64 | 7 | 91 |
| **128** | **3** | **64** | **3** | **32** | **3** | **90** |
| 128 | 3 | 64 | 5 | 32 | 7 | 97 |
| 128 | 5 | 64 | 5 | 32 | 5 | 92 |
| 128 | 7 | 64 | 5 | 32 | 3 | 91 |
| 128 | 7 | 64 | 7 | 32 | 7 | 93 |

Other layers were the same. The MAE ranged from 189 to 402 using one 1D CNN layer and ranged from 98 to 187 using two 1D CNN layers. The optimal architecture and parameters are consisted of three 1D CNN layers (MAE=90) with the least parameters compared to the other three 1D CNN architectures.

Figure 3 (A) shows that the input shape is (272, 21). The number of columns is the sum of heavy chain variable region length (145) and light chain variable region length (127). The rows came from one-hot encoding, including 20 amino acids and one gap. The input layer was connected with a 1D CNN layer using the activation function of the rectified linear unit (Relu). The number of filter and kernel sizes was 128 and 3, respectively. This layer was followed by a batch normalization layer connected to a dropout layer. The dropout rate was 0.5. The next was two 1D CNN layers and a max pooling layer. The nodes were then flattened to 1 dimension before connecting to a fully connected layer of size 32. Finally, the fully connected layer was connected to an output layer of size 1. The output layer was the SCM score of the antibody.

Figure 3 (B) plots the loss function of the best model over epochs. The MAE values for the training set dropped smoothly. The MAE values for the validation set initially increased and decreased rapidly until epoch 12. The optimal model was found at epoch 36. At this epoch, the MAE values for the training and validation sets were 86 and 90, respectively. After epoch 40, the model overfitted the training set, and the MAE value of the validation set started to climb.

Figure 3 (C) shows the scatter plot of the SCM scores and the DeepSCM scores on the test set. The linear correlation coefficient and MAE were 0.9 and 93.5, respectively. The MAE value was close to the training and validation sets, indicating a good model performance without significant overfitting. Furthermore, the MAE values were in the range of intrinsic fluctuation from MD simulations, as shown in the standard deviations in Figure 2 (B).

### 3.6 Applying DeepSCM to screen high concentration antibody viscosity

The performance of the DeepSCM as an effective surrogate model for the SCM model to screen high concentration antibody viscosity was assessed by 38 therapeutic antibodies from three different sources. The SCM scores correctly predicted the viscosity of these 38 antibodies. The criterion for high viscosity is > 30 cP at 150 mg/mL. When the SCM score is > 1000, the antibody is predicted to have high viscosity. Figure 4 plots the DeepSCM scores and viscosity at 150 mg/mL. Among the 38 antibodies, 18 were not in the dataset for the CNN model, and only one antibody was misclassified. Those in the dataset could be in the test sets that were not included for training. This result demonstrates that the DeepSCM model is a good surrogate model for the SCM model. Table 3 summarizes the SCM score, DeepSCM score, static SCM score for the 38 antibodies used in this test. Static SCM scores were calculated based on the homology models without MD simulations. There were only 21 mAbs correctly classified using the static SCM scores. The result indicates MD simulations were necessary for the SCM calculations, and the DeepSCM scores outperformed the static SCM scores.

**Table 3.**
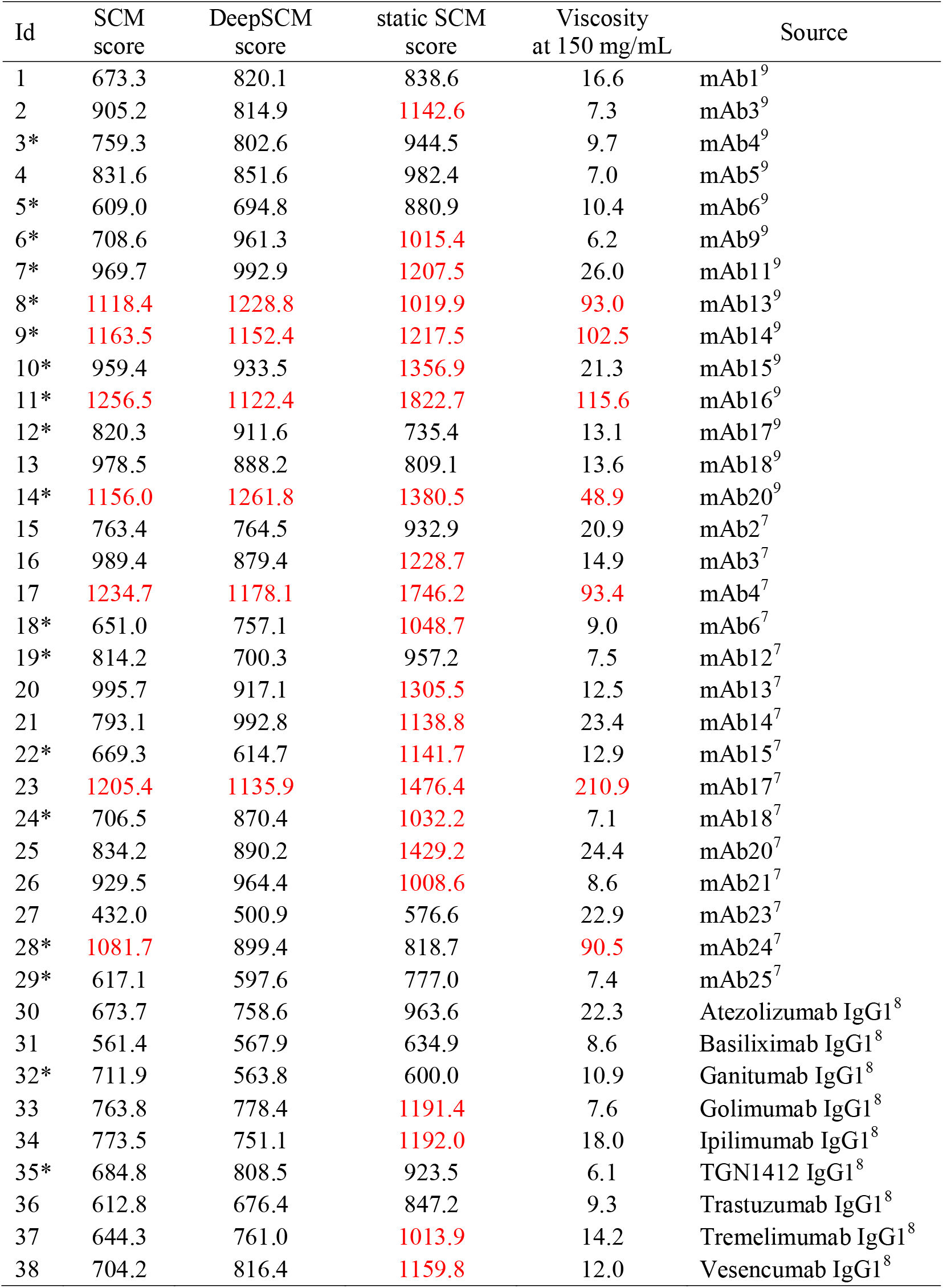
Summary of the DeepSCM score and the experimental viscosity at 150 mg/mL for the 38 mAbs. The SCM score is from the MD simulation, and the static SCM score is from the homology model. The red labels indicate predicted or experimental high viscosity. Their names in the source data are also listed. The asterisk signs indicate that the sequences are not in the dataset of this study.

| Id | SCM score | DeepSCM score | static SCM score | Viscosity at 150 mg/mL | Source |
| --- | --- | --- | --- | --- | --- |
| 1 | 673.3 | 820.1 | 838.6 | 16.6 | mAb1<sup>9</sup> |
| 2 | 905.2 | 814.9 | 1142.6 | 7.3 | mAb3<sup>9</sup> |
| 3* | 759.3 | 802.6 | 944.5 | 9.7 | mAb4<sup>9</sup> |
| 4 | 831.6 | 851.6 | 982.4 | 7.0 | mAb5<sup>9</sup> |
| 5* | 609.0 | 694.8 | 880.9 | 10.4 | mAb6<sup>9</sup> |
| 6* | 708.6 | 961.3 | 1015.4 | 6.2 | mAb9<sup>9</sup> |
| 7* | 969.7 | 992.9 | 1207.5 | 26.0 | mAb11<sup>9</sup> |
| 8* | 1118.4 | 1228.8 | 1019.9 | 93.0 | mAb13<sup>9</sup> |
| 9* | 1163.5 | 1152.4 | 1217.5 | 102.5 | mAb14<sup>9</sup> |
| 10* | 959.4 | 933.5 | 1356.9 | 21.3 | mAb15<sup>9</sup> |
| 11* | 1256.5 | 1122.4 | 1822.7 | 115.6 | mAb16<sup>9</sup> |
| 12* | 820.3 | 911.6 | 735.4 | 13.1 | mAb17<sup>9</sup> |
| 13 | 978.5 | 888.2 | 809.1 | 13.6 | mAb18<sup>9</sup> |
| 14* | 1156.0 | 1261.8 | 1380.5 | 48.9 | mAb20<sup>9</sup> |
| 15 | 763.4 | 764.5 | 932.9 | 20.9 | mAb2<sup>7</sup> |
| 16 | 989.4 | 879.4 | 1228.7 | 14.9 | mAb3<sup>7</sup> |
| 17 | 1234.7 | 1178.1 | 1746.2 | 93.4 | mAb4<sup>7</sup> |
| 18* | 651.0 | 757.1 | 1048.7 | 9.0 | mAb6<sup>7</sup> |
| 19* | 814.2 | 700.3 | 957.2 | 7.5 | mAb12<sup>7</sup> |
| 20 | 995.7 | 917.1 | 1305.5 | 12.5 | mAb13<sup>7</sup> |
| 21 | 793.1 | 992.8 | 1138.8 | 23.4 | mAb14<sup>7</sup> |
| 22* | 669.3 | 614.7 | 1141.7 | 12.9 | mAb15<sup>7</sup> |
| 23 | 1205.4 | 1135.9 | 1476.4 | 210.9 | mAb17<sup>7</sup> |
| 24* | 706.5 | 870.4 | 1032.2 | 7.1 | mAb18<sup>7</sup> |
| 25 | 834.2 | 890.2 | 1429.2 | 24.4 | mAb20<sup>7</sup> |
| 26 | 929.5 | 964.4 | 1008.6 | 8.6 | mAb21<sup>7</sup> |
| 27 | 432.0 | 500.9 | 576.6 | 22.9 | mAb23<sup>7</sup> |
| 28* | 1081.7 | 899.4 | 818.7 | 90.5 | mAb24<sup>7</sup> |
| 29* | 617.1 | 597.6 | 777.0 | 7.4 | mAb25<sup>7</sup> |
| 30 | 673.7 | 758.6 | 963.6 | 22.3 | Atezolizumab IgG1<sup>8</sup> |
| 31 | 561.4 | 567.9 | 634.9 | 8.6 | Basiliximab IgG1<sup>8</sup> |
| 32* | 711.9 | 563.8 | 600.0 | 10.9 | Ganitumab IgG1<sup>8</sup> |
| 33 | 763.8 | 778.4 | 1191.4 | 7.6 | Golimumab IgG1<sup>8</sup> |
| 34 | 773.5 | 751.1 | 1192.0 | 18.0 | Ipilimumab IgG1<sup>8</sup> |
| 35* | 684.8 | 808.5 | 923.5 | 6.1 | TGN1412 IgG1<sup>8</sup> |
| 36 | 612.8 | 676.4 | 847.2 | 9.3 | Trastuzumab IgG1<sup>8</sup> |
| 37 | 644.3 | 761.0 | 1013.9 | 14.2 | Tremelimumab IgG1<sup>8</sup> |
| 38 | 704.2 | 816.4 | 1159.8 | 12.0 | Vesencumab IgG1<sup>8</sup> |

**Figure 4.**
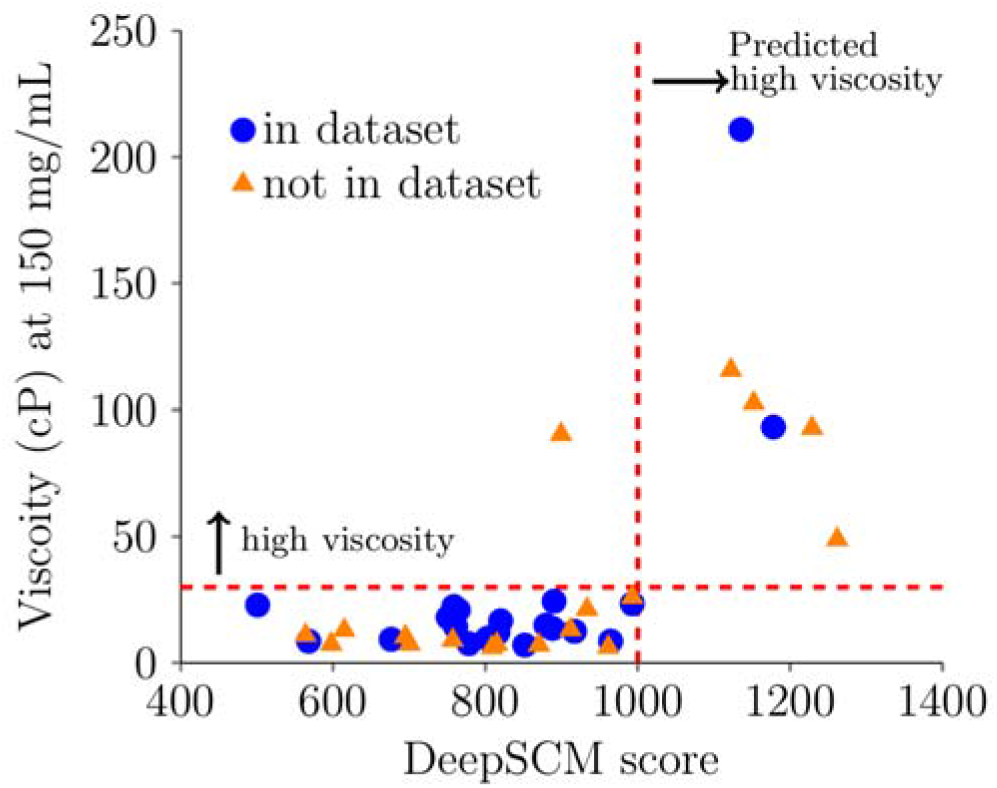
Experimental viscosity at 150 mg/mL with the DeepSCM score of 38 therapeutic antibodies. Blue circles are in the dataset of this study, either in training, validation, or test. Orange triangles are not in the dataset. The horizontal dashed line is the threshold value for high viscosity (30 cP), and the vertical dashed line is the threshold value for predicted high viscosity (1000).

### 3.7 Availability and implementation of the DeepSCM model

DeepSCM source code and pretrained parameters are freely available at https://github.com/Lailabcode/DeepSCM. Figure 5 shows the implementation flowchart. The FASTA files of the heavy chain and light chain Fv sequences need to be numbered by the ANARCI program using the IMGT definition in a CSV format. The seq_preprocessing.py program combines the numbered heavy chain and light chain CSV files to generate an input file. Currently, the preprocessing step ignores any insertions on the framework regions based on the IMGT definition. In addition, residues longer than the maximum length of each CDR region are excluded. The pred.py program takes the input file to calculate the DeepSCM scores.

**Figure 5.**
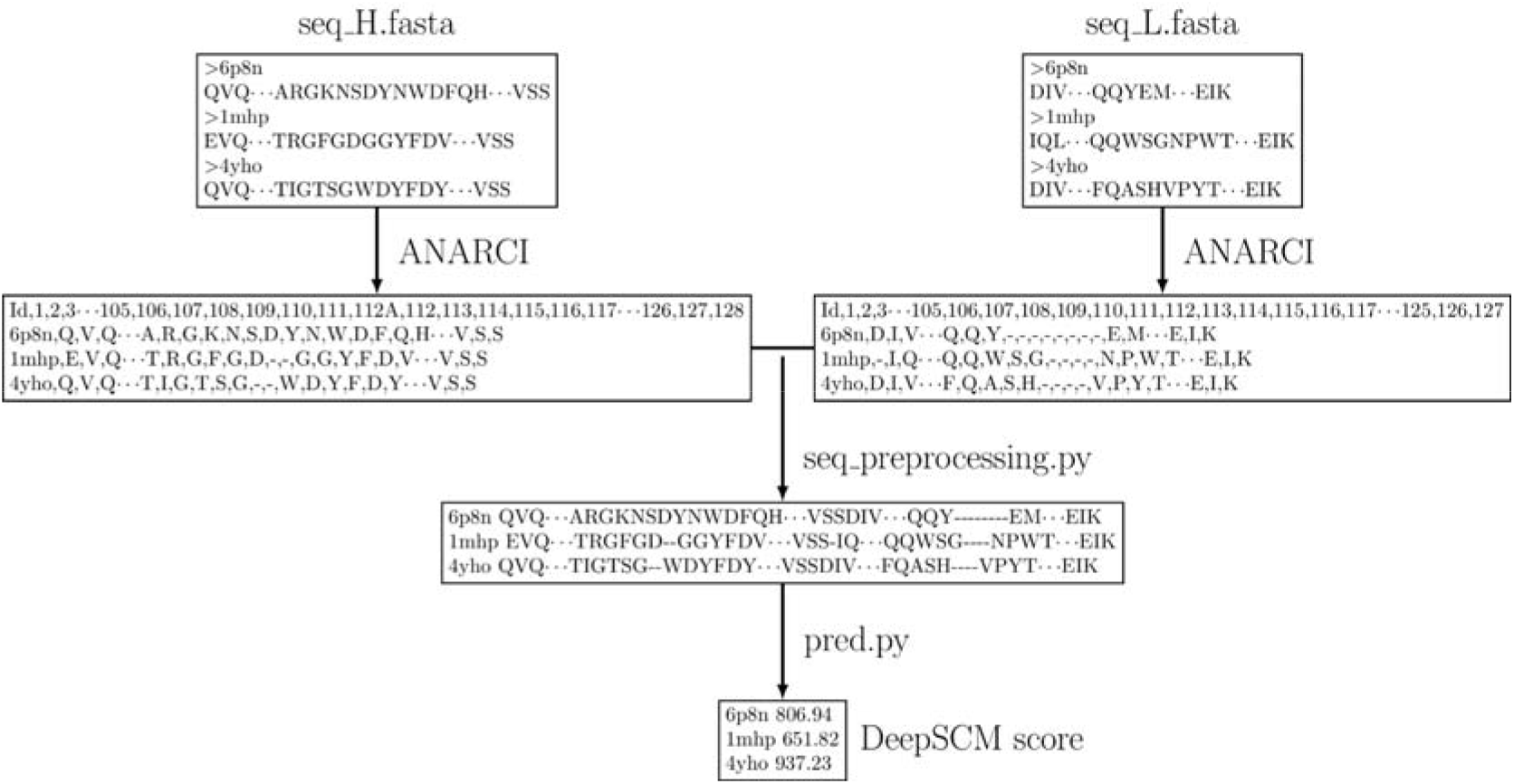
The flowchart of the DeepSCM program. The input files are FASTA files of heavy and light chains. The ANARCI program is used to number the heavy chain and the light chain antibody sequences in the IMGT scheme. The seq_preprocessing.py program converts the numbered sequences into DeepSCM input format. The pred.py will output the DeepSCM score.

## 4. Discussion

The advantages of the DeepSCM using CNN models over the SCM using MD simulations are twofold. The first advantage is speed. MD simulations for an antibody take several hours to days using modern supercomputers; however, CNN models take only a few seconds on personal computers. In addition, the CNN models require no homology models. The second advantage is reproducibility. MD simulations are stochastic by nature; therefore, the SCM scores vary slightly every run, even with the same input, making it challenging to transfer models with others. On the other hand, the sequence-based DeepSCM model guarantees exact reproducibility.

It remains a challenge to interpret the physical meaning of deep learning models. By developing a surrogate model for MD simulation, the interpretability of the CNN model is derived from the physically-based model. The accuracy of the DeepSCM model to predict therapeutic antibody viscosity also depends on the underlying assumption of the SCM model. The SCM model accounts for the surface exposed negative charges on the Fv region, a major driving force to induce high viscosity. However, other factors like aromatic rings and hydrophobic patches that could contribute to elevated viscosity are not included in the SCM model.^41^ Improving physical models to describe antibody viscosity behavior is still an outstanding problem.

The protocol described in this study to simulate many antibody sequences for deep learning training can be applied to other structural descriptors such as spatial aggregation propensity^42^ or solvent accessible surface area. In this work, only the antibody sequences from public databases were used for training. The training dataset can be augmented by a combinatorial design of different antibody regions to improve the deep learning model. In addition, biophysical properties from experiments such as melting temperature, retention time from hydrophobic interaction chromatography, and self-interaction from charge-stabilized self-interaction nanoparticle spectroscopy^43^ could be trained by the CNN model if combined with high throughput screening to generate larger datasets. These are potential future research topics. Deep learning paves a promising way for predicting antibody functions to facilitate drug design.

## 5. Conclusion

DeepSCM was developed as a surrogate model for MD simulation-based high concentration antibody viscosity screening tool. It was trained using high-throughput MD simulation results and 1D convolutional neural network architecture. DeepSCM enables viscosity screening for hundreds of antibody drug candidates using only antibody Fv sequences within a few seconds. This tool will facilitate early-stage drug development.

## Abbreviations

CDR: complementarity determining region
CDRH1: the first complementarity determining region of the heavy chain
CDRH2: the second complementarity determining region of the heavy chain
CDRH3: the third complementarity determining region of the heavy chain
CDRL1: the first complementarity determining region of the light chain
CDRL2: the second complementarity determining region of the light chain
CDRL3: the third complementarity determining region of the light chain
CG: coarse-grained
Fv: variable fragment
IgG1: immunoglobulin G1
IgG4P: immunoglobulin G4 S228P
mAbs: monoclonal antibodies
MD: molecular dynamics
SCM: spatial charge map
VH: heavy chain variable region
VL: light chain variable region

## Acknowledgments

The preclinical and clinical data were from a previous MIT-AstraZeneca collaboration. I thank Neil Mody and Austin Gallegos for generating the data. I thank Theresa Cloutier for providing insightful feedback on the manuscript. I thank the extreme science and engineering discovery environment (TG-CHM210013 and TG-CHM210016) for supporting computing resources.

